# Adapted protocol for *Saccharibacteria* co-cultivation: two new members join the club of Candidate Phyla radiation

**DOI:** 10.1101/2021.07.23.453610

**Authors:** Ahmad Ibrahim, Mohamad Maatouk, Andriamiharimamy Rajaonison, Rita Zgheib, Gabriel Haddad, Jacques Bou-Khalil, Didier Raoult, Fadi Bittar

## Abstract

The growing application of metagenomics to different ecological and microbiome niches in recent years has enhanced our knowledge of global microbial biodiversity. Among these abundant and widespread microbes, Candidate Phyla Radiation or CPR have been recognised as representing a large proportion of the microbial kingdom (> 26%). CPR are characterised by their obligate symbiotic or exo-parasitic activity with other microbial hosts, mainly bacteria. Currently, isolating CPR is still considered challenging for microbiologists. The idea of this study was to develop an adapted protocol for the co-culture of CPR with a suitable bacterial host. Based on various sputa, we tried to purify CPR (*Saccharibacteria* members) and to cultivate them with pure hosts. This protocol was monitored by real-time PCR quantification using a specific system for *Saccharibacteria* designed in this study, as well as by electron microscopy and sequencing. We succeeded in co-culturing and sequencing a complete genome of two new *Saccharibacteria* species: *Candidatus* Minimicrobia naudis and *Candidatus* Minimicrobia vallesae. In addition, we noticed a decrease in the Ct number of *Saccharibacteria*, and a significant multiplication through their physical association with *Schaalia odontolytica* strains in the enriched medium that we developed. This work may help bridge gaps in the genomic database by providing new CPR members and, in the future, their currently unknown characteristics may be revealed.

**IMPORTANCE:** In this study, the first real-time PCR system has been developed. This technique is able to quantify specifically *Saccharibacteria* members in any sample of interest in order to investigate their prevalence. In addition, another easy, specific and sensitive protocol has been developed to maintain the viability of *Saccharibacteria* cells in an enriched medium with their bacterial host. The use of this protocol subsequently facilitates studying the phenotypic characteristics of CPR and their physical interactions with bacterial species, as well as the sequencing of new genomes to improve the current database.

## INTRODUCTION

Over the past two decades, the fast progress of molecular methods and the intensive use of both total and targeted metagenomics (mainly 16S ribosomal RNA gene) have led to the recognition of new microorganisms which were not previously reported (1, 2). These recently-described microbes, which now represent a huge and diverse proportion of the microbial domain, are generally microorganisms that have not yet been cultured (1). Following each major discovery, and according to a recent classification based on whole genome content analyses, CPR are beginning to appear as a new division in the rhizome of life, independent from classical bacteria (3, 4). Since these microbes are not present in a pure cultivable state, their phenotypic characteristics remain incompletely defined (5). All known data are simply extracted from predictions based on bioinformatics analyses, which encourages microbiologists to culture them (1, 5). However, many difficulties limit their culture, such as slow growth/division, the need for specific metabolites in the final medium, and the growth inhibition by other dominant microorganisms or, inversely, the need for an obligatory association with another microorganism serving as a host in order to flourish (1, 2, 6, 7).

Recent studies on microbial diversity in human and environmental samples based on whole metagenomics analyses has made it possible to identify a new group of microorganisms that are not well recognised by the 16S rRNA gene, and which continue to be resistant to culture: the Candidate Phyla Radiation (also named CPR) (8). This group is comprised of more than 73 new Phyla, and represents a huge proportion (more than 26%) of the bacterial domain (2, 9, 10). Although CPR members present high inter-individual heterogeneity in genomic sequences, they do have certain common characteristics; they are morphologically small (100 to 300 nm), have a reduced genome size (usually less than 1 Mgb) (1), a high percentage of hypothetical proteins (11), and a single copy of 16S rRNA (8). Furthermore, CPR have a developed cell membrane close to that of Gram-positive bacteria (11), as well as limited and unknown/undetailed biosynthetic and metabolic capacities (12). In addition, they are enriched by proteins involved in cell-cell interactions, such as the presence of Pili belonging to the type IV secretion system (13). These proteins allow CPR members to be attached to their respective hosts, characterising their lifestyle, which appears to be either an exosymbiotic or exo-parasitic relationship (6, 7, 13).

Recently, it has been suggested that CPR co-evolved with bacteria (and not from bacteria), based on the distribution and diversity of their protein families (4, 11). Recent studies have shown that CPR are unable to synthesise nucleotides *de novo* and that they retain only the genes essential for their survival (11, 14). In fact, CPR seem to behave in a different, particular way (a non-traditional biological process), with their own ribosomal structures, and introns are present in their transfer RNA (tRNA) and 16S rRNA sequences (12). Analysis of the genomes available in the NCBI (National Centre for Biotechnology Information) database has led to the prediction of certain phenotypic characteristics unique to this group of microbes. These characteristics include their natural resistance to bacteriophage, despite the absence of the CRISPR viral defence in their genomes, which is due to the lack of viral receptors in their cell membrane (15), and the presence of different proteins involved in Quorum Sensing phenomena and cell-cell communication (16). None of these characteristics, however, have yet been confirmed *in vitro*.

*Saccharibacteria* or TM7, is the most studied CPR phylum and was named due to its sugar metabolism (17). Sequences belonging to this phylum have been systematically detected in several environmental and ecological samples, including soil, freshwater lakes, dolphin teeth, and termite guts (18, 19). In addition, metagenomics studies have shown that members of TM7 are also present in the human microbiome, including the intestinal, oral, urinary, cutaneous, blood and vaginal microbiota (11, 17, 20–22). Various studies have shown that *Saccharibacteria* members are associated with various human mucosal-related diseases, such as vaginosis, periodontitis and bowel disease (6, 20, 23).

To date, a few members of *Saccharibacteria* have been co-cultured with different Gram positive and negative bacterial hosts, most often *Schaalia odontolytica, Actinomyces* spp.*, Cellulosimicrobium cellulans, Lachnoanaerobaculum saburreum, Arachnia propionica* and *Leptotrichia* spp. (1, 2, 6, 24). Based on streptomycin resistance prediction, TM7x HMT-952 (also known as *Candidatus* Nanosynbacter lyticus) was the first TM7 strain to be cultivated and sequenced with its bacterial host in 2015 (6).

In order to expand our knowledge about this phylum, and to improve its phenotypic characterisation, culture is essential. Our aim in this study was to develop an easy and reproducible protocol for purifying strains belonging to *Saccharibacteria* species recovered from a human oral sample and to co-cultivate them with a mixture of *Schaalia odontolytica* strains.

## RESULTS

### 1 Specificity of the real time PCR system

The specificity of our designated qPCR system was confirmed using the collection of DNAs mentioned above. All bacterial and fungal DNA samples were negative, as well as the 25 stool samples which were negative on 580-F–1177R specific primers for *Saccharibacteria*. For greater accuracy, we tested 25 different sputum samples. All samples were positive by standard PCR and by our designated real-time PCR, with Ct values ranging between 17.02 and 23.57. In addition, the BLASTn analysis of the amplicons sequenced by Sanger shows that they all matched with different *Saccharibacteria* 23S rRNA genes. This system can amplify 126 base pair fragments of the 23S rRNA gene that serves as a specific marker for all *Saccharibacteria* spp.

### 2 Isolation and co-culture of *Saccharibacteria* species and quantification test

After checking that the two samples studied here were positive by specific real-time PCR for *Saccharibacteria* (similar Ct values were obtained for the two original samples tested (18.04 and 17.61 respectively), a seven-day period of enrichment, in TSB-BHI with hemin and vitamin K was initiated. Given that CPR members have a physically reduced corpuscle, they can pass through a 0.45-0.22 filter, allowing for efficient isolation of CPR cells for co-culturing and sequencing. In addition, we managed to concentrate *Saccharibacteria* cells in high quantities by ultracentrifugation (Figure 1). Most of the reads obtained by MiSeq-Illumina and GridION sequencing corresponded to *Saccharibacteria* sequences. After mixing the pellet with the 6 *S. odontolytica* strains, and due to the protocol steps, the Ct value of each sample was respectively 23.02 and 23.78). Co-culturing was then monitored by qPCR. In both samples, we noticed a significant decrease in Ct values after 48 hours of culture (21.07 and 21.24 respectively) (Figure 2). However, after this step and until the eighth day of culture, no significant variations in Ct values were observed. Ct values remained almost stable. The presence of *Saccharibacteria* cells at each step was also confirmed by electron microscopy (Hitachi TM4000 Plus and SU5000) following the presence of exosymbiotic coccus attached to several bacterial forms (Figure 3).

**Figure 1:**
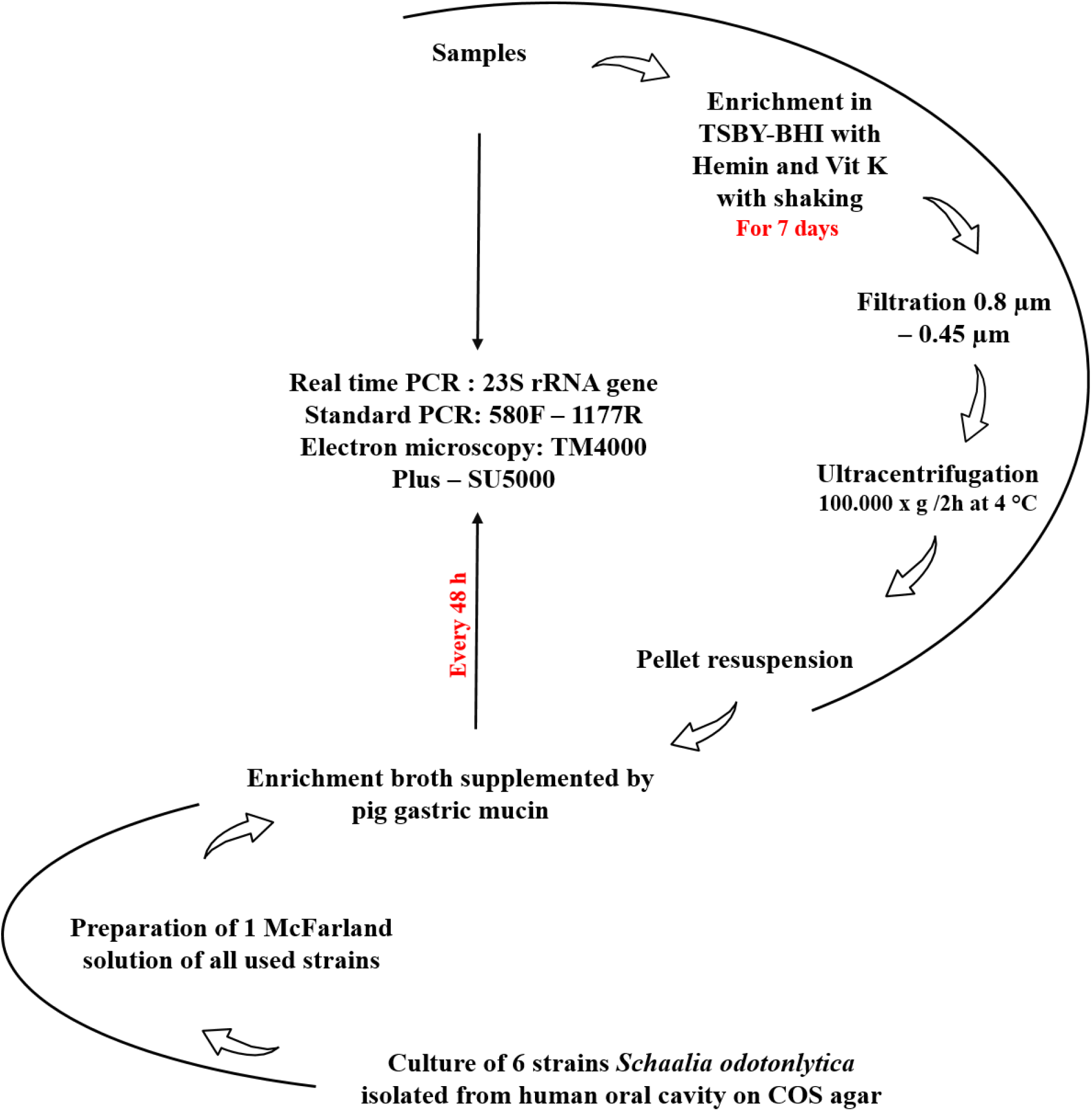
Summary of the *Saccharibacteria* co-culture protocol used in this study.

**Figure 2:**
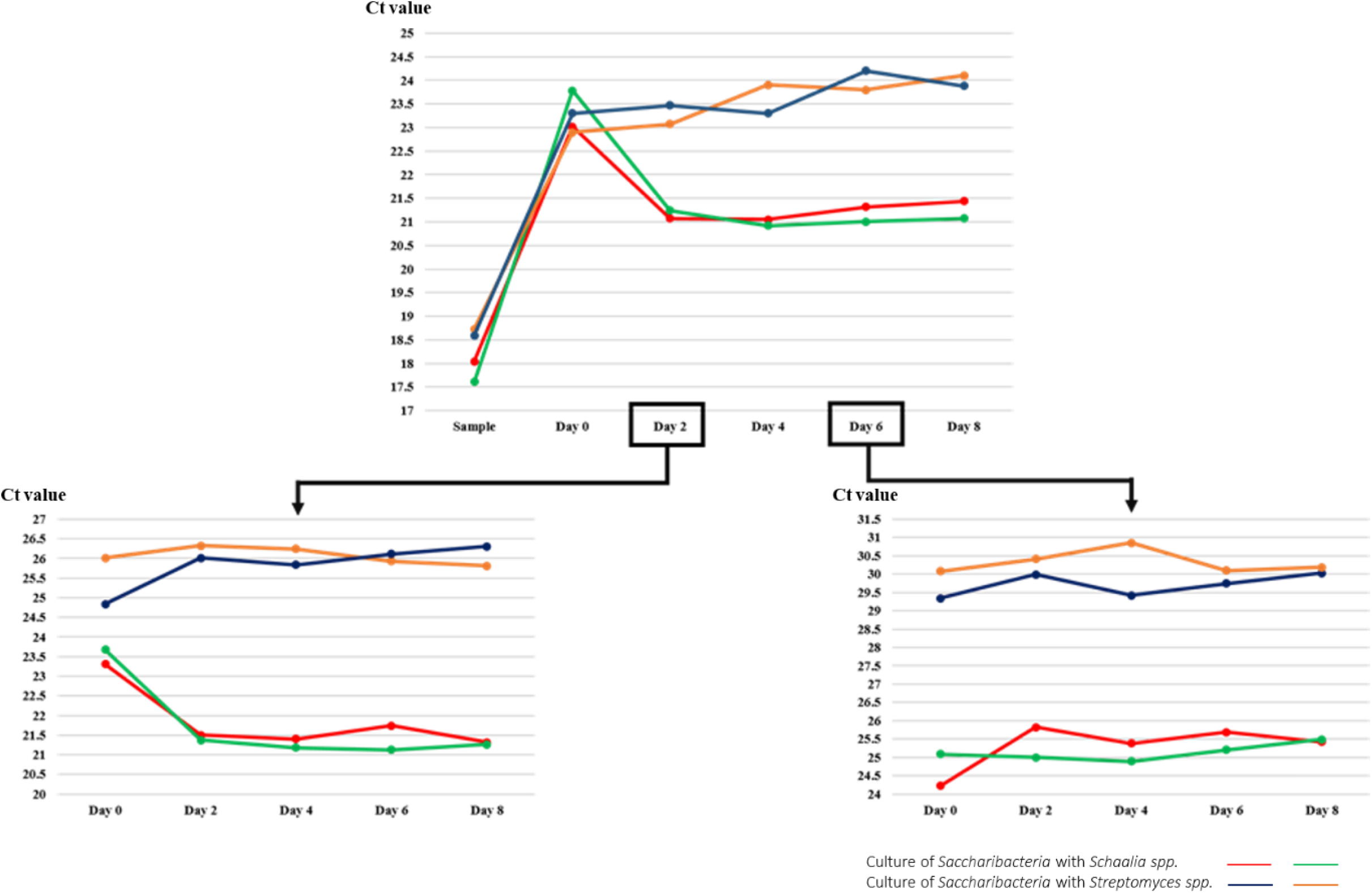
Graphic representation showing the Ct variation between each condition tested in this study. The co-culture of first sample with *Schaalia odontolytica* is represented in red, and the second is represented in green. For the co-culture with *Streptomyces* strains, the anaerobic conditions are indicated in blue and the aerobic condition in yellow.

**Figure 3:**
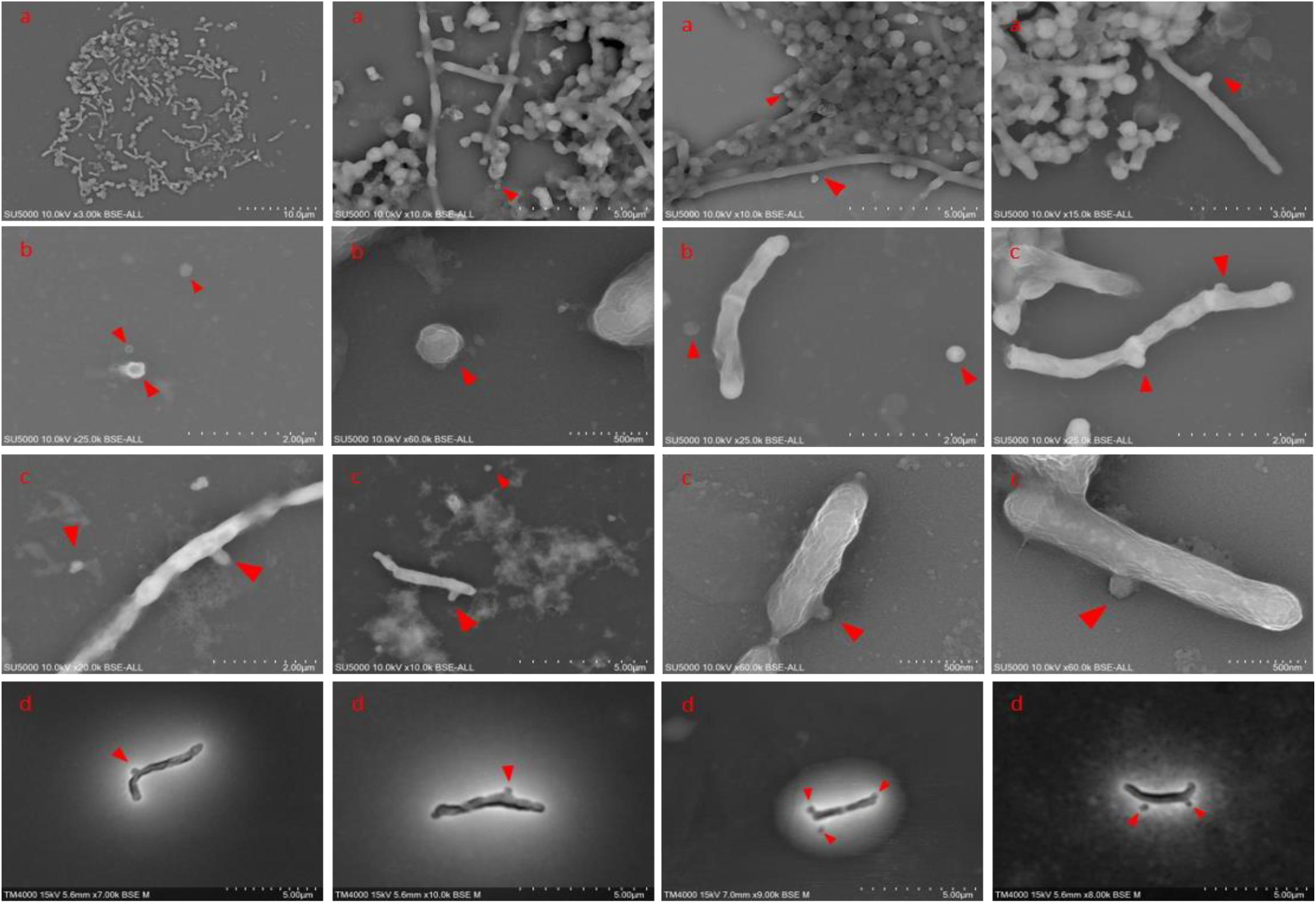
(a) Electron microscopy micrographs showing the presence of *Saccharibacteria* in the samples tested, (b) the *Saccharibacteria* cells detached from their host after filtration, (c) the physical association between purified *Saccharibacteria* cells with their new host (*Schaalia odontolytica*) using Hitachi – SU5000 and (d) using Hitachi TM4000 Plus.

To ensure that the nutrients were continuously renewed, a passage was performed on the second and sixth day of culture; 200 μl of the enrichment broth (containing *Saccharibacteria* cells and *S. odontolytica* strains) was mixed with 2.4 ml of initial medium supplemented with pig gastric mucin and incubated at 37°C in anaerobic conditions. Due to the dilution factor (200 μl in 2.4 ml), the Ct values were higher on day 0 of the passage (day 2 of the initial co-culture) in both samples (25.07 and 24.87 respectively) (Figure 2). We obtained comparable results: after only 48 hours of incubation, Ct values were also lower (23.61 for the first sample and 23.9 for the second). Conversely, we observed no multiplication of CPR following the passage made from the sixth day of the initial enrichment (Figure 2). This test confirms the viability of *Saccharibacteria* cells attached to *S. odontolytica* and the success of CPR co-culture using this protocol.

However, co-culturing of the pellets of a third sample (starting Ct= 18.92) with the three *Streptomyces* strains did not render similar results. The Ct values remained stable afterwards for eight days. Even the two passages did not increase the Ct values in aerobic and anaerobic conditions. Thus, the *Saccharibacteria* cells did not multiply following their association with this new bacterial host (Figure 2).

Finally, after 48 hours of co-culture, between 50 and 100 μl of each enrichment broth was deposited on COS medium, SHI supplemented with blood and mucin, and BHI supplemented with 10% sheep blood. Each anaerobically isolated colony was tested by qPCR. Our qPCR system could not identify positive colonies. For greater precision, a standard PCR test was performed, and all colonies were negative for *Saccharibacteria*. The MALDI-TOF-MS (Matrix-Assisted Laser Desorption Ionisation Time-of-Flight Mass Spectrometry) test identified the most isolated colonies as *S. odontolytica* (or *A. odontolyticus*) / *Streptococcus oralis* with a high score (>1.9). This score indicates the absence of foreign proteins (such as *Saccharibacteria* proteins) in each colony that can affect the spectra related to each known bacteria.

### 3 *Saccharibacteria* cell imaging by electron microscopy

Each initial sample was observed using electron microscopy (Hitachi TM4000 Plus and SU5000). We noticed a strong presence of biofilm, and a lot of coccus microbes attached to the external surface of several bacterial forms (bacilli and cocci). The size of these particles ranged from 100 and 400 nm, which corresponds to the described size of CPR members (Figure 3).

However, following the filtration/centrifugation of the initial enrichment step, we were able to observe single and detached cocci forms, with no association with any bacterial host (Figure 3). The size of these particles is similar to those observed in the original samples, and much smaller than the known cocci bacteria (*Staphylococcus* spp*., Streptococcus* spp. for example). These observations, along with the molecular results, confirm that *Saccharibacteria* cells were well separated from their bacterial hosts (Figure 3).

Finally, a microscopic slide for each host strain used was viewed using the two electron microscopes; we were unable to detect any form with a size similar to that of CPR cells. However, round-shape cells (1 to 2/ bacterial cell) appeared on the surface of these strains on the second day of their co-culture with the “purified” *Saccharibacteria* spp. (Figure 3), and single *Saccharibacteria* and *S. odontolytica* cells continued to be observed. Hence, a physical association between purified *Saccharibacteria* and its host appeared. There were, therefore, bacteria that did not harbour CPR, and other bacteria that were carriers of a maximum of one or two *Saccharibacteria*. The observations on day 4 and day 6 showed the same results.

### 4 Genomic sequencing and description

For each DNA sample, the total of Illumina and Nanopore reads were mapped against the *Saccharibacteria* reference genome (TM7x) using the CLC genomics 7 server. The filtration protocol, combined with the pre-treatment extraction allowed us to cover the entire TM7x genome (100%) in each DNA sample. Using long range PCR, we obtained two complete genomes representing two new *Saccharibacteria* species. The first genome (named *Candidatus* Minimicrobia naudis) has a length of 708,351 bp with 43.9% G+C content. It has 1,324 protein coding genes that include 792 hypothetical proteins (59.81%). Similarly, the second sequenced genome (*Candidatus* Minimicrobia vallesae) has a length of 706,973 bp and a 43.7% of G+C content. 48.97% of its protein coding genes (n= 1,017) correspond to hypothetical proteins (Supplementary data: Table S1). In addition, according to the proteomic analysis, 719 and 618 protein-coding genes of *Candidatus* M. naudis and *Candidatus* M. vallesae, respectively, were assigned to COGs categories (Supplementary data: Figure S1, Table S2). We did not detect any proteins belonging to the following COGs categories: B, Q, W, X, Y & Z. A graphic circular map for each genome is presented in Supplementary data: Figure S2. Genomic comparison between our two genomes and TM7x (as a reference genome) using Easyfig v-2.2.5 is presented in Figure 4. In addition, recent studies have shown the presence of introns in the tRNA of CPR (12). Here, we identified one tRNA/genome which contains intronic sequence: Gly CCC for *Candidatus* M. naudis and Thr TGT for *Candidatus* M. vallesae (Figure 5). We did not find any NRPS/PKS clusters nor IS sequences in either genome. With regard to antimicrobial resistance screening, *Candidatus* M. naudis was resistant to mupirocin, glycopeptide, tetracycline and oxazolidinone. Likewise, we found resistance genes for glycopeptide, tetracycline, oxazolidinone and MLS in the *Candidatus* M. vallesae genome (25). Finally, we also found type II, IV and VI Pili secretion systems in both genomes, and type I Pili in *Candidatus* M. vallesae only.

**Figure 4:**
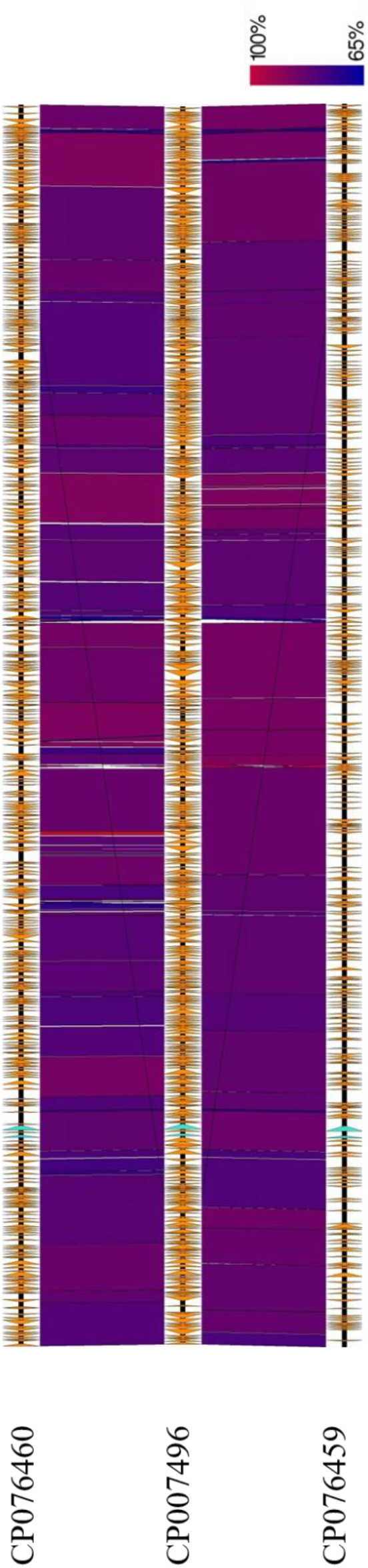
(a) Graphic representation showing the genomic comparison between *Candidatus* Minimicrobia vallesae, (b) the TM7x reference genome and (c) and *Candidatus* Minimicrobia naudis. This representation was generated using the Easyfig v 2.2.5 online tool.

**Figure 5:**
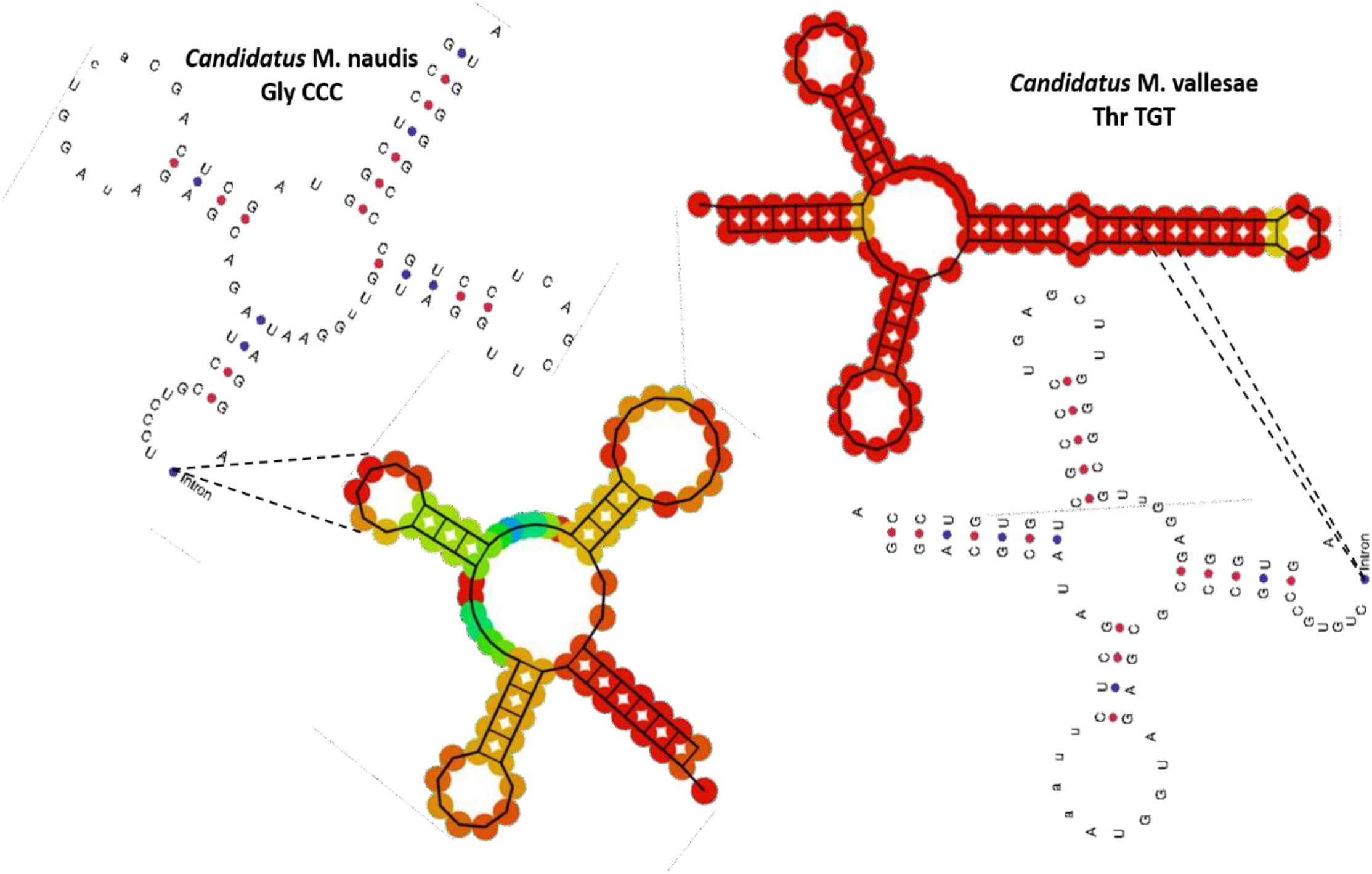
Two-dimensional representation of tRNA with intronic sequences, detected in each genome.

For taxogenomic classification, the phylogenetic trees based on 16S rRNA, and the whole genome sequences show that our two new *Minimicrobia* species belong to the superphylum *Saccharibacteria*, with all the CPR tested phyla (Figure 6). In addition, the analyses of 16S ribosomal RNA, as previously described, show that our two new species belong to the clade G1 of *Saccharibacteria* oral species (26, 27). The maximum OrthoANI value was 84.2412% for *Candidatus* M. naudis with TM7 - ASM569739v1, 84.0275% for *Candidatus* M. vallesae with TM7-ASM80362v1 and 90.7707% between them (Supplementary data: Figure S3). Likewise, digital DNA-DNA hybridisation showed that our described genomes had the highest values (26.01% for *Candidatus* M. naudis, 25.3% for *Candidatus M. vallesae*) with *Candidatus Saccharibacteria* bacterium oral taxon 955 - ASM1020192v1 and TM7-ASM569739v1, respectively. The percentage between them was 41.7% (39.2–44.2 confidence interval). According to these values, we defined *Candidatus* M. naudis and *Candidatus* M. vallesae as two new CPR species belonging to the *Saccharibacteria* phylum.

**Figure 6:**
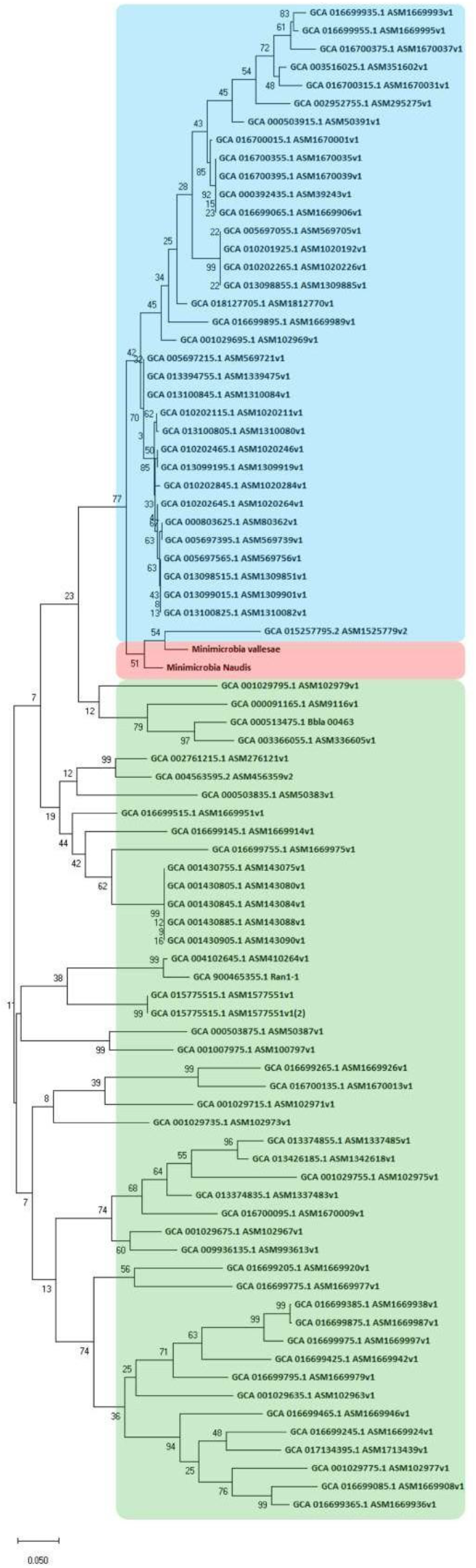
Unrooted phylogenetic tree shows the analyses of the 16S ribosomal RNA gene of all available *Saccharibacteria* complete genomes (marked in blue), *Candidatus* Minimicrobia naudis (marked in red), *Candidatus* Minimicrobia vallesae (marked in red) and all available non-*Saccharibacteria* CPR complete genomes (marked in green). This tree was generated using MegaX.

According to the taxonomic affiliation of each *Saccharibacteria* sequence, their origins were determined. The evolutionary history of each genome is presented here based on all genomic sequences belonging to the repertoire of coding genes. We obtained a particular mosaicism for both *Candidatus* M. naudis and *Candidatus* M. vallesae (4), similar to one another and comparable to that of the reference genome (Figure 7).

**Figure 7:**
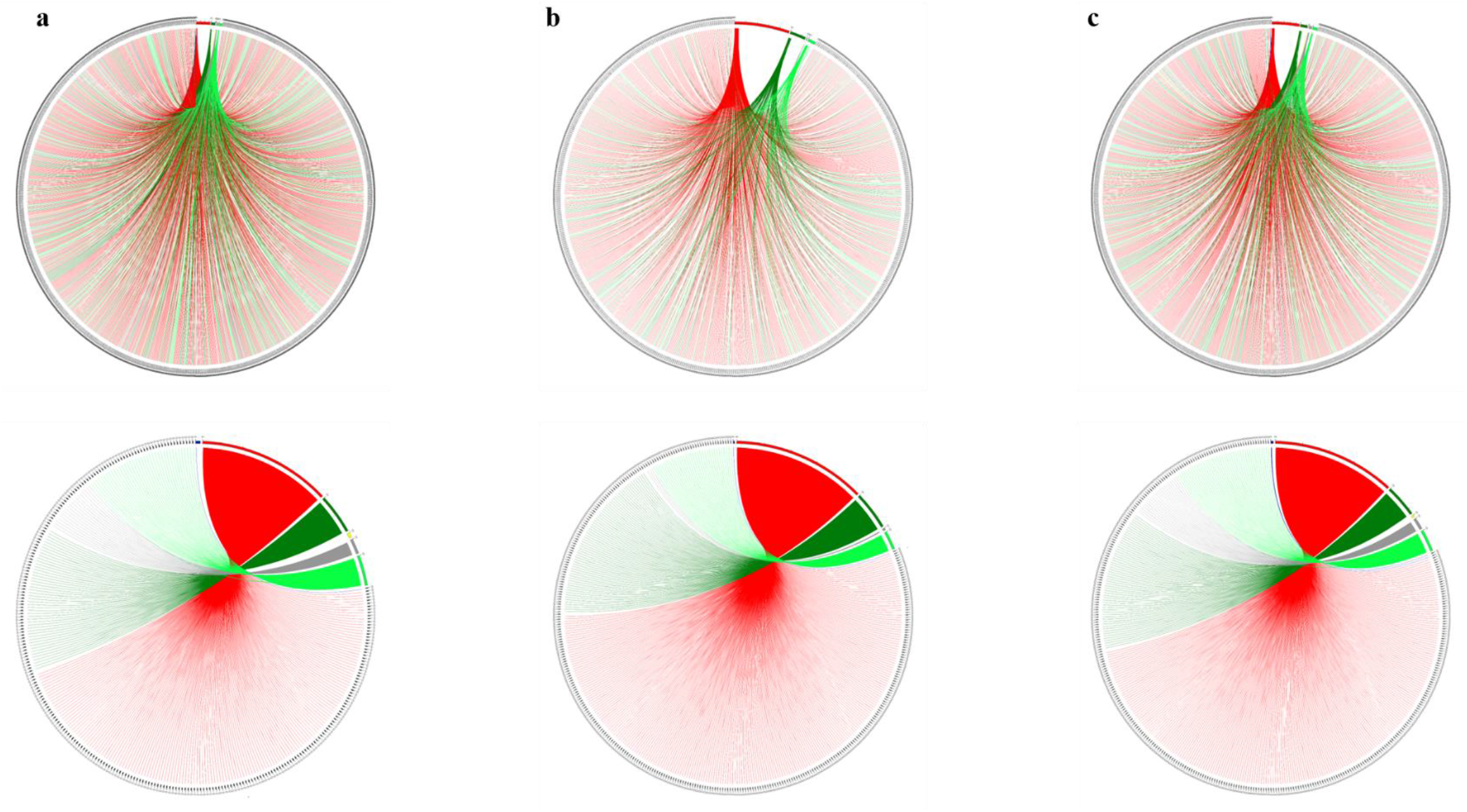
Rhizomal illustration presenting the mosaicism of each used genome: (a) *Candidatus* Minimicrobia naudis, (c) *Candidatus* Minimicrobia vallesae and (b) TM7x reference genomes. Each protein encoding gene is represented by a curve, coloured according to its origin: bacterial origin in red, CPR non-*Saccharibacteria* phylum origin in dark green, *Saccharibacteria* phylum origin in light-green, Eukaryotic origin in yellow, archaeal origin in dark-blue, and ORFans in grey. In the first line, each curve represents a protein encoding gene, arranged in the figure by order. For the second line, protein encoding genes belonging to the same origin are arranged together. Figures were performed using the circos tool

For each genome, we found a prevalence of sequences of bacterial and CPR origins (45.6% and 42.6%, respectively, for *Candidatus* M. naudis and 48.4% and 45.4% for *Candidatus* M. vallesae, respectively). Among the sequences of CPR origin, a large percentage is unique to the phylum *Saccharibacteria* (an average of 31% in each genome). However, we also detected some eukaryotic and archaean sequences in each genome (0.32% / 0.24% respectively, for *Candidatus M.* naudis and 0.16% / 0.3% for *Candidatus* M. vallesae, respectively) (Figure 7).

## DISCUSSION

The oral microbiota is known as the most complex human microbiota. It has been estimated that it may contain more than 775 microbial species (22). In addition, following the initial inclusion of CPR in the tree of life, different metagenomics studies have shown that the *Saccharibacteria* phylum is very abundant in humans and, more precisely, in the oral cavity (17). Therefore, this co-culture protocol was mainly tested on sputum samples.

The quantification and viability of *Saccharibacteria* has been tested by standard PCR in a number of studies (1, 6, 24, 28). Different sets of primers targeting the 16S rRNA have been identified as universal for this phylum (28). According to these results, *Saccharibacteria* was considered viable if the PCR was still positive after five passages (1). This method increases the risk of false positive results by amplifying DNA for dead microorganisms and/or mis-quantification. Here, we developed a real time qPCR system which was, for the first time, specific for *Saccharibacteria* phylum, enabling us to detect and quantify this phylum in any interesting samples. In addition, this specificity was re-confirmed by selecting all additional complete genomes available in the NCBI database between 1 December 2019 and 1 June 2021. This system was able to amplify 34/35 tested genomes (specificity = 97.12%). Following our results, very low Ct values were obtained from fresh sputa, confirming their abundance in the oral microbiota (17). Secondly, *Saccharibacteria* members have not yet been cultivated in pure culture. Their identification on agar media or by MALDI-TOF MS is currently impossible. The use of this system, followed by metagenomics analysis, therefore enable this phylum to be screened in any sample, and in the future may lead to greater precision regarding their prevalence in humans and in environmental samples.

In this study, in line with several others (1, 2, 6), we confirmed that *Saccharibacteria* cells (CPR cells in general) can detach themselves from their natural host bacteria following continuous agitation. They can then adapt with another host to multiply (1). Following a co-culture of *S. odontolytica* strains with purified *Saccharibacteria* cells, Ct values decreased after two days, which explains their persistence and viability in liquid media. However, Ct values remained stable between days 2 and 8. It could, therefore, be suggested that the nutrients needed by CPR cells have already been consumed and/or the metabolic and nutrient transport between the host and the guest has entered a standby stage, hence we were unable to detect further multiplication.

Nutritional supplementation of this complex (renewal of enrichment passage at day 2) restored these activities. Two criteria should therefore be considered to keep CPR at the multiplying stage: having a suitable host and a well-renewed enriched medium. In addition, and as suggested by He *et al.*, CPR accompanies *Schaalia* spp. in stable long-term infections due to the adaptation and rapid evolution of its host (6). Moreover, it is thought that, on day 6 of culture, the CPR were dead, and only the DNA of the dead cells was amplified. Therefore, we failed to decrease the Ct value after a passage from the sixth day of initial culture. The protocol optimised in this study therefore guarantees highly protection and easy purification of the CPR and ensures very sensitive monitoring of their viability by electron microscopy and qPCR. It also provided the *Saccharibacteria* with an enriched nutrient complex, especially with the addition of pig gastric mucin during the host infection stage. This protocol could be used to search for other bacterial hosts not yet described for CPR.

It is known that the physical size of the CPR is between 100 and 300 nm, so we limited the filtration here to 0.45 µm, to avoid losing a quantity of CPR between 0.22 µm and 0.3 µm. Therefore, our metagenomics analyses of the filtrate showed some contaminations of sequences belonging to the *Streptococcus* and *Veillonella* species which passed through the filters (29). However, most of the reads still correspond to the phylum TM7 - *Saccharibacteria*.

Furthermore, we were unable to isolate a positive colony as demonstrated by our real-time PCR system. Following a deposit of the starting sample and the filtrate mixed with *Schaalia* spp., all colonies were negative in real time PCR and electron microscopy. A recent study showed that the use of reverse genomics methods was successful in producing *Saccharibacteria* positive colonies (24). This method is based on a target antibody that only picks up *Saccharibacteria* with their hosts (24). In our assay, other microorganisms were able to pass through the 0.45 µm filtrate. We suggest that the requirement of *Saccharibacteria*, and/or their fragility by the presence of other microorganisms in the filtrate (*Streptococcus oralis* for example), prevented their multiplication on a solid medium, even though several enriched media were tried (COS, supplemented BHI and SHI agar). It would, therefore, be interesting to find universal epitopes, common to all known *Saccharibacteria* rather than based on one or two genomes, to facilitate their solid culture and sorting them using flow cytometry.

It is known that *Saccharibacteria* members interact with *S. odontolytica* to multiply in an exo-symbiotic (or exo-parasitic) relationship, in stable long-term infections between these two microorganisms. Furthermore, different studies have suggested that *Saccharibacteria* spp. can adapt with other bacteria, such as *Arachinia* spp. for example (1). Here, the infection of *Streptomyces* spp. by purified *Saccharibacteria* cells was not successful in terms of their multiplication, indicating that the association between these microorganisms is not appropriate to a nutrient transfer from the host bacterium to the CPR cells. Hence, *Streptomyces* cannot be considered as one of the hosts of identified *Saccharibacteria* species. Finally, this protocol extends the described diversity of CPR to date. It enabled us to recover two new species belonging to the phylum *Saccharibacteria*. Both species are unique, and they are of similar size to those described in the literature, but with very divergent sequences (maximum OrthoANI and DDH values are very low). In addition, we found a tRNA with intronic sequences in each genome, which has recently been described in CPR genomes (12).

Concerning their origin, the presence of archaeal/eukaryotic sequences suggests the presence of an interaction between these microorganisms in their shared niche (4, 30, 31). The mosaic structure of CPR in general gives them a unique characteristic, comparable to one another and different from other microbial domains (4).

## MATERIALS AND METHODS

### 1 Sample collection and ethics statement

Twenty-eight sputum samples were collected at La Timone University Hospital (AP-HM, Assistance Publique-Hôpitaux Marseille) from routine laboratory diagnostics. Research analyses were only performed on surplus samples, once laboratory diagnostic procedures had been initiated. The patients were informed that their samples may be used for research purposes and retained the right to oppose to this use. Given that this study did not involve specific collection of samples or use medical/personal data from patients, and according to French law (the Jardé’s law), neither institutional ethical approval nor individual patient consent was required for this non-invasive study (Loi no 2012–300 du 5 mars 2012 and Décret no 2016–1537 du 16 novembre 2016 published in the ‘Journal Officiel de la République Française’).

Each 2 ml sample was diluted in 1 ml of transport medium composed of 0.1g MgCl_2_, 0.2 g KH_2_PO_4_, 1.15 g NaCl, 1g Na_2_HP_4_, 1g ascorbic acid, 1g uric acid and 1g Glutathione per 1 litre of deionised water; pH= 7.5). All tested samples were stored in anaerobic conditions.

### 2 Isolation of *Saccharibacteria* spp. and culture conditions

In a hemoculture tube, we diluted 1 ml of each sputum sample in 39 ml enriched broth: (per 1,000 mL TSB (Tryptic soy broth): 37g BHI (Brain Heart Infusion), 10g yeast extract, 10 mg Hemin and 50 μl Vitamin K pH final = 7, (BioMérieux, Marcy-l’Etoile, France) at 37 °C and in an atmosphere of 85% N_2_, 10% CO_2_ and 5% H_2._ Each culture was performed in anaerobic chamber (Coy), for seven days with agitation (300 rpm) to separate the *Saccharibacteria* cells present from their bacterial hosts. After seven days of enrichment and agitation, the broth was filtered at 0.8 μm and 0.45 μm respectively, to eliminate big particles and associated cultured bacteria. For greater cell concentration, an ultracentrifugation of 100.000 x g was then performed for two hours at 4°C. The pellet (which was sometimes invisible) was resuspended in 2.5 ml of the enrichment broth mentioned above, supplemented with 2.5g/l of pig gastric mucin.

In addition, we prepared a 1 McFarland solution of six *Schaalia odontolytica* strains (previously known as *Actinomyces odotonlyticus*), isolated from a human oral cavity, in physiological water. For each resuspended pellet, 200 μl was used for molecular biology analyses and the remaining quantity was cultured with 0.1 ml of *Schaalia odontolytica* strains solution for seven days in a Hungate tube with no agitation, under the same anaerobic conditions described above. After 48 hours of culture, 50 to 100 μl of each enrichment broth was deposited on COS agar, SHI and BHI agar (BioMérieux, Marcy-l’Etoile, France) supplemented by 10% sheep blood and 2.5 pig gastric mucin each in anaerobic conditions (Figure 1). The same culture protocol described above was also tested on other samples by mixing the filtrate with 1 McFarland of three

*Streptomyces* spp. strains (*Streptomyces cattleya* DSM 46488, *Streptomyces massiliensis*, *Streptomyces rochei*), isolated from the human gut, separately in aerobic and anaerobic conditions (Figure 1).

### 3 *Saccharibacteria* viability testing

To evaluate *Saccharibacteria* co-culture, we designated a real time PCR system for quantification. To do so, we selected all *Saccharibacteria* complete genomes available in the NCBI on 1 December 2020. Based on the conserved ribosomal genes, a multiple alignment of 23S ribosomal was performed to determine the conserved zones. We consequently selected SacchariF: GGCTTATAGCGCCCAATAG as a forward primer, SacchariR: CGGATATAAACCGAACTGTC as a reverse primer, and SacchariP: 6-FAM-CATAGACGGCGCTGTTTGGCAC-TAMRA as a TaqMan probe.

The specificity of this system was confirmed *in silico* by BLASTn against the nr database, and i*n vitro* against a variety of 50 bacterial species, 70 *Candida* strains (32) and 25 stool samples which had previously tested negative with the specific *Saccharibacteria* standard PCR (580-F – 1177-R) (33).

To improve the extraction of *Saccharibacteria* DNA, several pre-treatments were performed on each tested sample and culture condition. For deglycosylation, each 180 μl sample/*Saccharibacteria* co-culture was treated with Endo Hf Kit P0703L (New England Biolabs, Evry, France): 3 μl of each reagent was added to the sample and incubated for one hour at room temperature, then one hour at 37°C. We then added 10 μl lysozyme for two hours, 10 μl proteinase K for 12 hours at 56°C, followed by one-minute disruption with glass powder using Fast-Prep. We used the EZ1 biorobot (Qiagen BioRobot EZ1-, Tokyo, Japan) for the automated extraction, using the EZ1 DNA tissue kit (EZ1 DNA, Qiagen, Hilden, Germany) and the bacterial protocol card. Each sample of DNA extracted was diluted in 50 μl solution. A PCR quantification test was then performed on each sample before culture, and every 48 hours after infecting *Schaalia odontolytica* strains / *Streptomyces* spp. strains with purified *Saccharibacteria* cells. For this purpose, we used the CFX-96 device connected to TM-BioRad using TaqMan technology (Figure 1). The qPCR reactions were carried out according to the following protocol: two minutes of incubation at 50°C, 15 minutes of activation at 95°C, followed by 40 cycles of five seconds at 95°C and 30 seconds at 60°C for DNA amplification, then a final step at 45°C for 30 seconds. We prepared each qPCR mixture in 20 μl total volume containing 10 μl of QuantiTect Primers Assays, 2 μl of sterile water, 1 μl of each primer, 1 μl of probe, and 5 μl of each DNA (32). In addition, to confirm the qPCR specificity, each amplicon was sequenced using the Sanger method and analysed by BLASTn against the nr database.

### 4 Bacterial and CPR imaging

All specimens or samples were fixed in 2.5% glutaraldehyde solution and were deposited by cyto-centrifugation on cytospin slides, followed by staining with a 1% PTA (Phosphotungstic acid) aqueous solution (pH =7) for three minutes. All samples were then sputtered with a 10 nm thick Platinum layer to reduce charging of the imaged samples.

For image acquisition, we first used Hitachi’s TM4000 Plus tabletop SEM, approximately 60 cm in height and 33 cm wide to evaluate bacterial structure. We used the Backscatter Electron (BSE) as a detector. The voltage of acceleration was 10 kV and magnifications varied from 250 X to 7,000 X. Using the same accelerating voltage, we then used Hitachi’s SU5000 SEM for the higher resolution and magnifications. Magnifications varied from 5,000 X to 15,000 X. The evacuation time after loading specimens into the SEM Chamber was less than two minutes. All co-cultures of samples / *Saccharibacteria* cells were acquired using the same acquisition settings regarding magnification, intensity and voltage mode. Here, each microbial form presenting a cocci shape (coccus), and a physical size between 100 and 400 nm, outside or attached to a bacterium was considered as a CPR cell.

### 5 Next-generation sequencing

Extracted DNA was sequenced using two different methods, firstly on the MiSeq (Illumina Inc, San Diego, CA, USA) using the Nextra XT DNA sample prep kit (Illumina), with the paired end strategy. The tagmentation step fragmented and tagged each extracted DNA to prepare the paired-end library. A limited PCR amplification (12 cycles) was then performed to complete the tag adapters and to introduce dual-index barcodes. DNA was then purified on AMPure XP beads (Beckman Coulter Inc, Fullerton, CA, USA). In addition, according to the Nextera XT protocol (Illumina), all libraries were normalised on specific beads. We then pooled all libraries into one library for DNA sequencing on MiSeq. The pooled single strand library was loaded onto the reagent cartridge and then onto the instrument along with the flow cell. Automated cluster generation and paired end sequencing with dual index reads were performed in a single 39-hour run in 2×250-bp.

The Oxford Nanopore method was then performed on 1D genomic DNA sequencing for the GridION device using the SQK-LSK109 kit. A library was constructed from 1 µg genomic DNA without fragmentation and end repair. Adapters were ligated to both ends of the genomic DNA. After purification on AMPure XP beads (Beckman Coulter Inc, Fullerton, CA, USA), the library was quantified by a Qubit assay with the high sensitivity kit (Life technologies, Carlsbad, CA, USA). We detected active pores for sequencing and the WIMP workflow was chosen for live bioinformatic analyses.

### 6 Genomic description

For each sample/purified *Saccharibacteria* DNA sequence, the quality of each Illumina and Oxford Nanopore read was checked by FastQC, and trimmed using trimmomatic version 0.36.6. We merged all the reads that corresponded to a given sample (this protocol was applied to one sputum, and then confirmed on a second). Each group of reads was mapped against the reference *Saccharibacteria* genome (*Candidatus* Nanosynbacter lyticus, available on NCBI under accession number: ASM80362v1) using CLC Genomics Workbench v.7. We used the default parameters except for the length fraction (reduced to 0.3) and the similarity fraction (reduced to 0.5). Mapped reads were assembled using SPAdes software, version 3.13.0 (34) using the default options. For this step, we only kept contigs with a minimum size of 400 bp. Each contig was then analysed by BLASTn against the nr database and we only kept contigs which matched with sequences corresponding to *Saccharibacteria* spp. All selected fasta sequences were then mapped against TM7x - ASM80362v1, using the same criteria mentioned above to generate the sequenced *Saccharibacteria* genome, with no contamination by any bacterial/eukaryotic sequences.

To complete our sequenced genomes (to fill in the gaps) we designed primers around each gap to perform long range PCR. Each PCR product (amplicon) was then sequenced using the Oxford Nanopore method and mapped against the contig to link them. These genomes were deposited in the GenBank as complete genomes under accession numbers CP076459 and CP076460. Then, coding and non-coding genes, hypothetical proteins, CDS, and rRNA were predicted using Prokka (35). tRNA genes were predicted by tRNA SCAN SE, using the default option and all available sequences sources (36). Proteomes were predicted with BLASTp (e-value of 0.001, minimum coverage and identity of 70% and 30% respectively) against the cluster of orthologous groups database (37). Antibiotic resistance genes were then predicted by Abricate (mass screening of contigs for antimicrobial and virulence genes) using ARG-ANNOT (antimicrobial resistance gene annotation) and ResFinder as databases (38). Similarly, we looked for the presence of NRPS-PKS using BLASTp against the NRPPUR database (39). In addition, in order to detect lateral sequence transfers between our species and their host (presence of transposon/integron), a screening for IS sequences was performed by BLASTn and BLASTp against the ISfinder online tool (40). *Saccharibacteria* members are known to have protein secretion systems (Pili) which attach onto the external membrane of their host. For this purpose, we screened our assembled genomes against the MacSyDB/TXSSdb online database (41) to detect all proteins secretion systems which were presented. An additional genomic comparison between our genomes and TM7x reference genome was then performed using Easyfig v.2.2.5 (42).

To determine the mosaicism and evolutionary history of each genome, we constructed a representative rhizome that showed the genetic exchange between our sequenced *Saccharibacteria* spp. and the other organisms (4). For this purpose, a BLASTp for each coding gene was performed against the NCBI protein database. Any protein which did not match with any sequence was considered as ORFans. The remaining best HITs were selected based on the following criteria: minimum identity and coverage of 20% and 30% respectively, and maximum e-value of 0.001, as previously described (4, 43). Rhizome representations were then constructed using the circos software (44).

For taxonomic characterisation, we selected for comparison all CPR complete genomes which were available on the NCBI at 1 June 2020 (n=81). A multiple alignment of 16S rRNA sequences was performed using MUSCLE software, and curated alignments were then used for the construction of a phylogenetic tree using the maximum likelihood (ML) method, with 1,000 bootstrap replicates, using nearest-neighbour-interchange (NNI) with the Jones-Taylor-Thornton (JTT) model. Tree were constructed using MEGA-X software. In addition, the degree of genomic similarity between all selected genomes was estimated using OrthoANI software. We also used the Genome-to-Genome Distance Calculator Web Service to calculate the digital DNA-DNA hybridisation (dDDH) value with confidence intervals according to recommended parameters, as a previously described (45).

## DATA AVAILABILITY

*Candidatus* Minimicrobia naudis & *Candidatus* Minimicrobia vallesae genomes were deposited in the NCBI-GenBank under accession numbers CP076459 and CP076460, respectively.

## ACKNOWLEDGMENTS

We thank Ludivine Brechard, Olivia Ardizzoni, Vincent Bossi, and Madeleine Carrara (Institut Hospitalo-Universitaire Méditerranée-Infection, Marseille 13385, France) for their technical help in sequencing. We would like to also thank Dr Linda Hadjadj and Dr Mouna Hamel for their committed assistance in the laboratory. We thank the Hitachi team of Japan (Hitachi High-Technologies Corporation, Science & Medical Systems Business Group 24-14, Nishi-shimbashi 1-chome, Minato-ku, Tokyo 105-8717 Japan) for the collaborative study conducted together with IHU Méditerranée Infection, and for the installation of the TM4000 Plus and SU5000 microscopes at the IHU Méditerranée Infection facility.

F.B., D.R., and A.I. designed the study. A.I. and M.M. wrote the manuscript. F.B., D.R. and J.B.K revised the manuscript. A.I. and M.M. performed the microbiological analyses. A.I., A.R. and R.Z. performed the bioinformatics experiments, A.I., G.H and J.B.K. performed the electron microscopy imaging. All authors have read and approved the final manuscript.

## FUNDING INFORMATIONS

This work was supported by the French Government under the « Investissements d’avenir » (Investments for the Future) program managed by the Agence Nationale de la Recherche (ANR,fr: National Agency for Research), (reference: Méditerranée Infection 10-IAHU-03). This work was supported by Région Provence-Alpes-Côte d’Azur and European funding (FEDER (Fonds européen de développement régional) PRIMMI (Plateformes de Recherche et d’Innovation Mutualisées Méditerranée Infection)). In addition, collaborative research conducted by IHU Méditerranée Infection and the Hitachi High-Tech Corporation is funded by the Hitachi High-Tech Corporation.

## SUPPLEMENTAL MATERIAL

**Figure S1:** Graphic representation showing the distribution of functional classes of predicted genes according to the clusters of orthologous groups (COGs) of proteins of *Candidatus* Minimicrobia naudis and *Candidatus* Minimicrobia vallesae, among other *Saccharibacteria* complete genomes.

**Figure S2:** Graphic circular map of *Candidatus* Minimicrobia naudis and *Candidatus* Minimicrobia vallesae complete genomes. Maps were generating using the CG view online tool.

**Figure S3:** Heat map generated with OrthoANI values calculated using the OAT software between *Candidatus* Minimicrobia naudis and *Candidatus* Minimicrobia vallesae and all CPR complete genomes available on NCBI at 1 June 2021.

**Table S1:** Genomic characteristics of *Candidatus* Minimicrobia naudis and *Candidatus* Minimicrobia vallesae.

**Table S2:** Distribution of functional classes of predicted genes according to the clusters of orthologous groups (COGs) of proteins of *Candidatus* Minimicrobia naudis and *Candidatus* Minimicrobia vallesae, among other Saccharibacteria complete genomes.

